# Grip and grasp: lizard claw inspired robotic manipulators

**DOI:** 10.1101/2025.10.22.683985

**Authors:** Hyeon Lee, Kate Manges Douglas, Asyiah Bray, Andrea Rummel, Parvez Alam

## Abstract

This paper considers the selection of a lizard claw to apply as an end effector on a bioinspired robotic manipulator, to improve grasping efficiency. Our work draws on morphological parameters that influence mechanical grip. By considering geometrical ratios that are built on parameters such as the arm-to-tip angle, the claw inner radius, and the slip distance, our research identifies optimal parameters that influence grip strength and stability. We demonstrate that these parameters, when combined to form two specific indices (the geometric mean, *I*_*GM*_, and the arithmetic mean of deviations, *I*_*AMD*_), provide a basis for more extensive comparisons between the different claw morphologies researched. The claw of *Crotaphytus collaris* is deduced as being the most suitable candidate for application as a bioinspired end effector in a robotic manipulator used for grasping objects, as it closely aligns with our optimal design criteria. To practically demonstrate gripping and grasping, we use a lightweight ping pong ball as well as larger and heavier objects such as an abalone shell, boxing mitts and a printer spool. Our robotic manipulator integrates dual grasping strategies. These include pinching for delicate objects and scooping for larger, irregular objects, thus enhancing its versatility across different environments and for different applications. Overall, this paper offers valuable new insights into biomimetic manipulator design using claws as end effectors, highlighting the importance of lizard claw selection based on morphology.

## 1. Introduction

Grasping in robotics is often viewed from the perspective of climbing [1–3]. This is because lizards are regularly observed performing grasping motions during climbing, while clinging, and as an aid to locomotion across topographically irregular surfaces [4–6]. The ability to grip is often enhanced by the presence of a claw [4,7,8] and while this may seem fundamentally obvious, detailed technical evidence to support this hypothesis has only really started appearing in the literature over recent decades. While smaller lizards may effectively use van der Waals forces to adhere to inclined and even, over-vertical substrates [9,10], these animals have claws, which benefit gripping when the surfaces are rough [7,8], i.e., when there is something to cling to. Certain morphological features are understood to be of potential benefit to climbing and gripping. Non-arboreal anoles, for example, have shorter claws and a smaller claw base-height than arboreal anoles [11]. The curvature of claws correlates with grip on smooth surfaces, while claw height correlates with grip on rough surfaces [12]. Tip sharpness affects grip through puncture [13,14], and various undimensionalised aspect ratios such as claw height to length, claw height to width, and claw curvature to tip angle, are parameters through which claws can be differentiated based on grip strength [4]. These and other morphological features in lizard claws have been a source of inspiration in the development of climbing robots. This includes the application of forward protruding claws from digits in robotic feet [15], the application of variable claw interaction angles with the substrate in climbing robots [16], the use of paired claws to increase grip strength [17], and a general use of curved claws to improve grip to rough substrates [18–20].

As discussed, the impetus for applied research on claw-inspired climbing technologies has grown substantially over the last decade [1,21]. There are also new technologies that focus on mimicking digit-claw *functionalities* to enhance other grip-grasp technologies, such as robotic manipulators, where the aim is to lift or hold onto an object [22–24]. Examples include manipulators that enable landing on thin structures such as branches [25,26], those that comprise actuated paired claws designed to collect objects in search and rescue operations [27], claws attached to manipulators that mimic interdigit adduction and abduction [28], and manipulators desgined as adaptive landing gears in unmanned aerial vehicles, versatile enough to land on uneven terrain, to grab objects, and to perch on cylindrical structures [29]. While there are several studies that mimic digit-claw functionalities, there are none that use true claw geometries. This is understandable as engineering design typically seeks to simplify for manufacturing simplicity, as well as to ensure safety through predictability. Nevertheless, with an almost exponential surge in additive manufacturing capabilities and tooling over the last decade, we are at a stage where we can and should consider using the true geometry of animal claws as this will help us ascertain optimal claw geometries for use in robotics. These can of course be simplified if needed, but we still need to ensure that our search space when designing claws as end effectors in manipulators includes analyses on real claw geometries. This paper begins that search, using a range of different lizard claws. Our work begins by testing and analysing the effects of claw geometry on grip properties and behaviours, after which we select and demonstrate the use of the optimal claw as an end effector applied to a tri-digit manipulator.

## 2. Results

### 2.1. Grip Testing

The grip test unit was connected to an Instron 3369 (Instron Inc., High Wycombe, Buckinghamshire, UK) to measure the force and displacement values of each claw type against a ping pong ball, aligning with similar sized and shaped gripping objects to those previously researched [30]. The grip test unit comprises an arm base, three arms positions at 120^*{*°^ relative to each other, and an upscaled, 3D printed claw attached to the base of each arm. Testing begins at the south pole of the ball and is continued until the claws slip from the ball. The claw grip unit is shown in Figure 1(a) (specifically here: *C. collaris*), where a sequence of still frames can be seen to the point of slip during a single test run. We repeated the test for each claw type eight times (n = 8). Representative (median) force-displacement curves are shown in Figure 1(b) for each of the claw types. Claw types with the higher median grip forces are found in the species: *C. texanus, C. collaris, B. vattatus, U. notata, T. hoehnelii*, and *S. arenicolus (forelimb)*. On observing the tests, we note that claw tests from the following species were very smooth and tended to follow the curvature of the ball: *P. modestum, S. arenicolus (hind), Salvator merianae, V. salvator (fore)* and *V. salvator (hind)*. As a result, the force-displacement curves for these claw types are parabola-like, indicating the work done to reach a maximum force is not released abruptly through slip. Species where an abrupt slip was observed include *B. vittatus, C. collaris, S. arenicolus (fore)* and *C. texanus*, each of which display an initial high stiffness region of the force-displacement curve, followed by a sharp reduction in load carrying capacity caused by slip. The force-displacement curves of claws from two species, *T. hoehnelii* and *U. notata*, plateau after reaching a force maximum. This was a result of the claws pressing into and deforming the ball across its equatorial planes. A final observation relates to claws from the species *C. giganteus*, which was unable to interface with the ball due to both its very low curvature and its angle relative to that of the ball.

**Figure 1.**
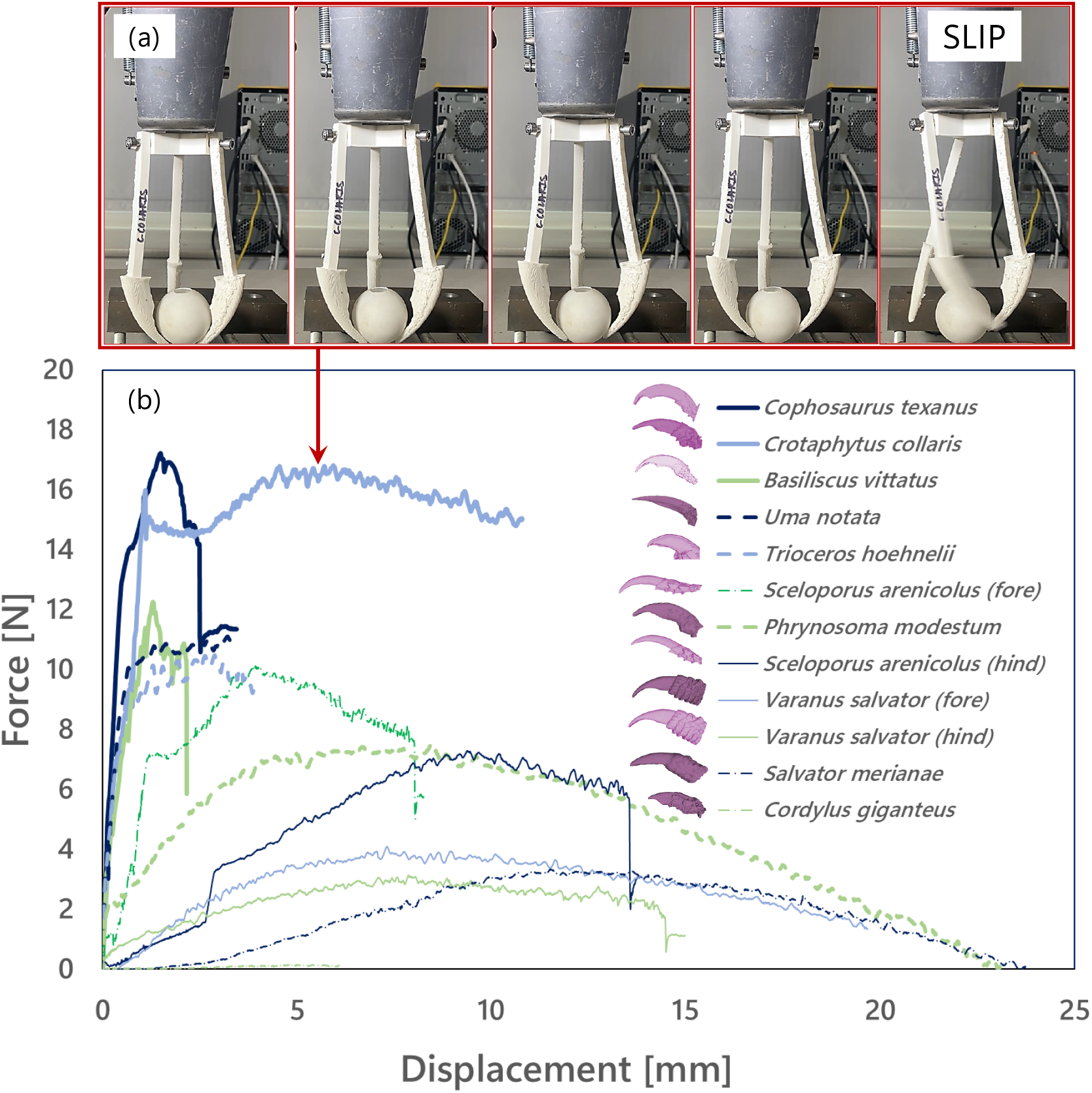
(a) sequence of still frames showing the grip test unit using claws from *C. collaris* to drag across the surface of the ping pong ball (with slip being included in the final frame) and (b) representative (median) force-displacement curves for each of the claw types tested with the grip test unit. In the figure legend, the species are ordered highest to lowest with respect to maximum force in these representative, and the claws shown adjacent to each name are not to scale.

Box and whisker plots shown in Figure 2(a) provide detail on the distribution of forces per sample set (claw type), while in Figure 2(b), the maximum displacements per sample set (claw type) are shown. Here, n = 8 per sample set. Tables providing numerical details on each, including standard deviations, standard errors and coefficients of variation, are provided as Electronic Supplementary Tables. In these plots, we note that *C. texanus, C. collaris, B. vattatus, U. notata, T. hoehnelii*, and *S. arenicolus (forelimb)*, while having higher mean and median force values than *P. moduestum, S. arenicolus hind), V. salvator (fore), V. salvator (hind), S. merianae* and *C. giganteus*, also tend to have larger distributions of force, indicating lower levels of consistency in terms of gripping (with the exceptions of *B. vittatus* and *S. arenicolus (fore)*). The reverse is noted in the displacement plot (with the exception of *U. notata* and *C. collaris*).

**Figure 2.**
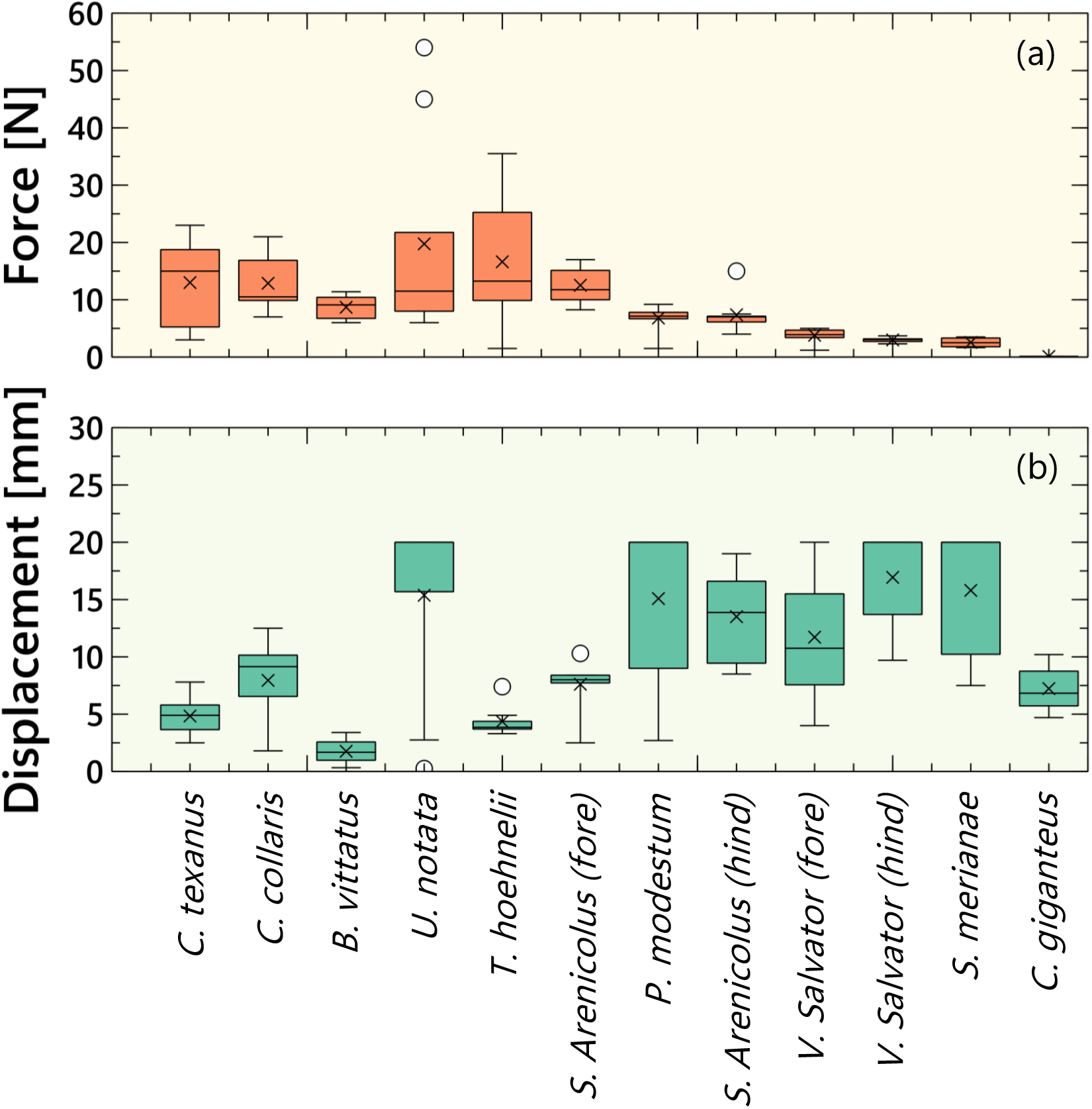
Box and whisker plots showing (a) the maximum forces reached and (b) the maximum displacement reached in each sample set (claw type), where n = 8 in each sample set.

### 2.2. Claw selection

Our prototype tri-digit robotic manipulator will use one claw type from a selection of several different lizard types. We use three separate criteria for claw selection. Each criterion is based on the dimensions and shape of a ping pong ball that we use as the ‘gripped substrate’ during testing. The first criterion considers general meshing of the ‘gripper’ against the ‘gripped’. We base this on simple angular alignment between the claw as attached to its arm, ∠_*AE*_. This conceptualisation is built on the understanding that should a rectilinear element be placed around a ball, it will form a right-angled square since the ball has an aspect ratio of 1:1. The angle of the south pole of a ball when placed against a flat vertical surface, ∠_*ball*_, is 135^°^. As such, to mesh perfectly with the ball using the ratio of angularity as a primary metric, *r*_1_, a robotic arm will also need to be at an angle of 135^°^ from the end effector (claw), Figure 3(a) and Equation 1. The closer *r*_1_ is to unity, the better the meshing of the gripper to the gripped substrate. The second criterion measures the closeness of the claw curvature to the ball and is also expressed as a ratio, *r*_2_, Equation 2. We base this on the understanding that optimal meshing is achieved when an estimated radius of curvature of a claw perfectly matches that of the ball, the radius of which is 20 mm. An *r*_2_ value of unity indicates perfect meshing, and meshing is considered to become progressively more inferior, the further this ratio diverges from unity. The concept is illustrated in Figure 3(b). While the first two criteria focus solely on geometry and meshing, the third criterion is measured from the vertical displacement of the claw tip relative to the south pole of the ball, Figure 3(c). An ideal claw grip will start slipping, *d*_*s*_, at the point of inflection of the ball relative to its south pole and its orthogonal. This will commence at a vertical displacement, *d*_*v*_, of one half of the ball radius, *r*_*ball*_, relative to the initial position of claw contact closest to the claw tip (the south pole of the ball). Here, 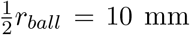 and the third ratio, *r*_3_, is shown in Equation 3. Here, 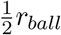 is considered as an optimal position from which slip begins, as we wish to maximise leverage and grip efficiency, while concurrently enabling consistent hinge and release motion. This design constraint is thence important as it directly impacts the predictability of robotic operational performance, precision, and adaptability in tasks involving gripping and manipulation.

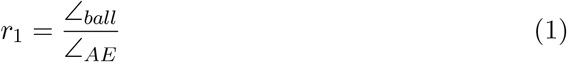

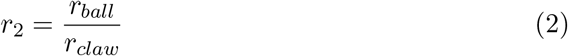

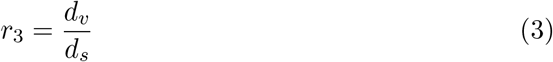

**Figure 3.**
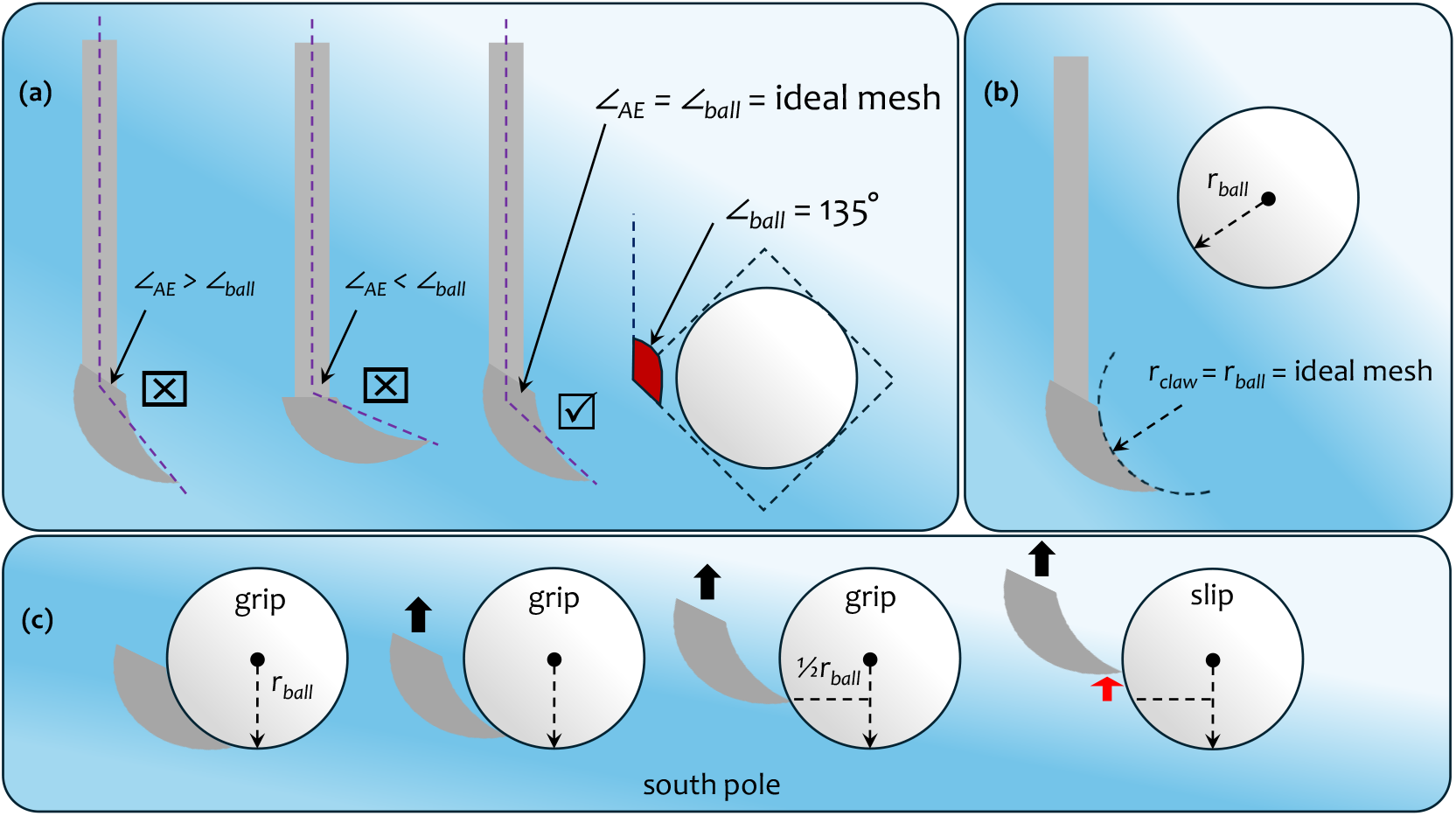
(a) representation of non-ideal and ideal angles between the arm and the end effector (claw) ∠_*AE*_ (b) representation of the ideal radius of curvature between the claw and the ball and (c) representation of the when grip between the ball and the claw will naturally decrease due to the gradient change between the two.

While each of the ratios described above, *r*_1_-*r*_3_, can act as standalone design criteria, these ratios can also be used to create two indices to further gauge the most optimal lizard claw geometry for our robotic manipulator. The first index is the Geometric Mean, *I*_*GM*_. This is a composite index that uses the three ratios as shown in Equation 4. An index value of 1 means all ratios are perfect; values deviating from 1 indicate how far the set is from the ideal.

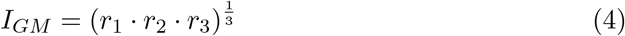

The second index is an Arithmetic Mean of Deviations, *I*_*AMD*_, which computes the absolute average of the individual ratios, as shown in Equation 5. This value will be maximised at 1 (when all ratios are 1) and decreases as ratios deviate from 1.

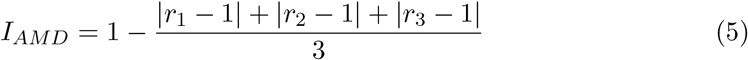

Figure 4 shows that the mean force, *F*, scales against the parameter ∠_*AE*_ using a linear regression model, *F* = *a*(∠_*AE*_) + *b*, where *a* and *b* are constants. Pearson’s correlation is a useful statistical measure as it enables an understanding of the strength and direction of a linear relationship between two continuous variables. It provides a single value, the Pearson correlation coefficient, *r*, which ranges from −1 to +1, which indicates how closely the data points fall to a straight line. Here, Pearson’s correlation is defined according to Equation 6, where 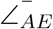 and 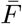 are the mean values of ∠_*AE*_ and *F*, respectively, *n* represents the number of samples, and 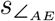 and *s*_*F*_ are the standard deviations of ∠_*AE*_ and *F*, respectively. Using Pearson’s correlation coefficient, we notice the negative linear relationship between *F* and ∠_*AE*_ is relatively high, *r* = *−*0.81, suggesting that as the independent variable ∠_*AE*_ increases, the dependent variable *F* tends to decrease markedly. The coefficient of determination (*R*^2^) for the linear regression fit is 0.65, indicating that 65% of the variance in the dependent variable (*F*) is explained by the model. Figure 5(a) compares each of *r*_1_-*r*_3_ in a choropleth map.

**Figure 4.**
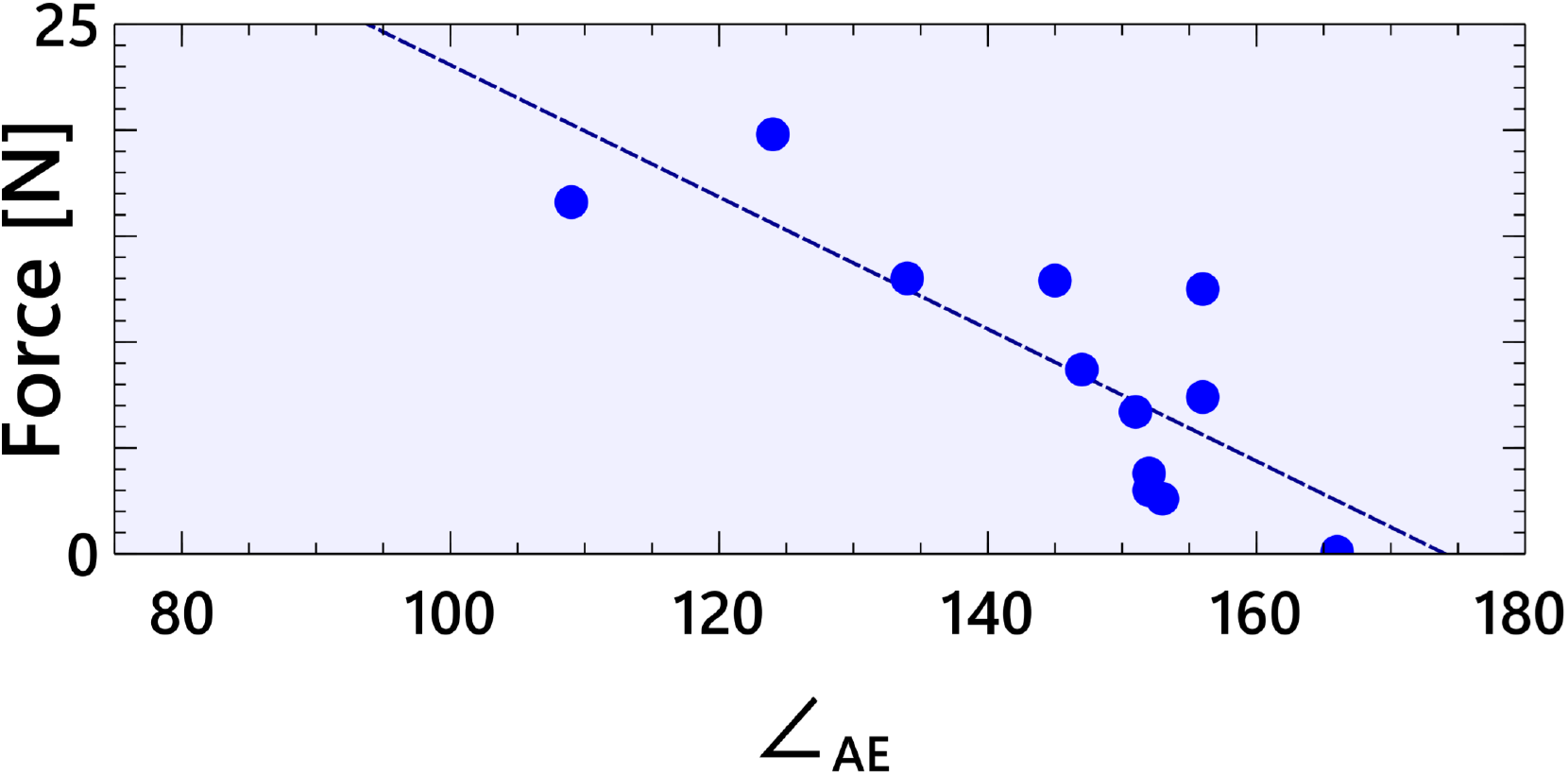
Mean force plotted against ∠_*AE*_ for each claw type with a linear regression fit applied: *r* = *−*0.81 and *R*^2^ = 0.65.

**Figure 5.**
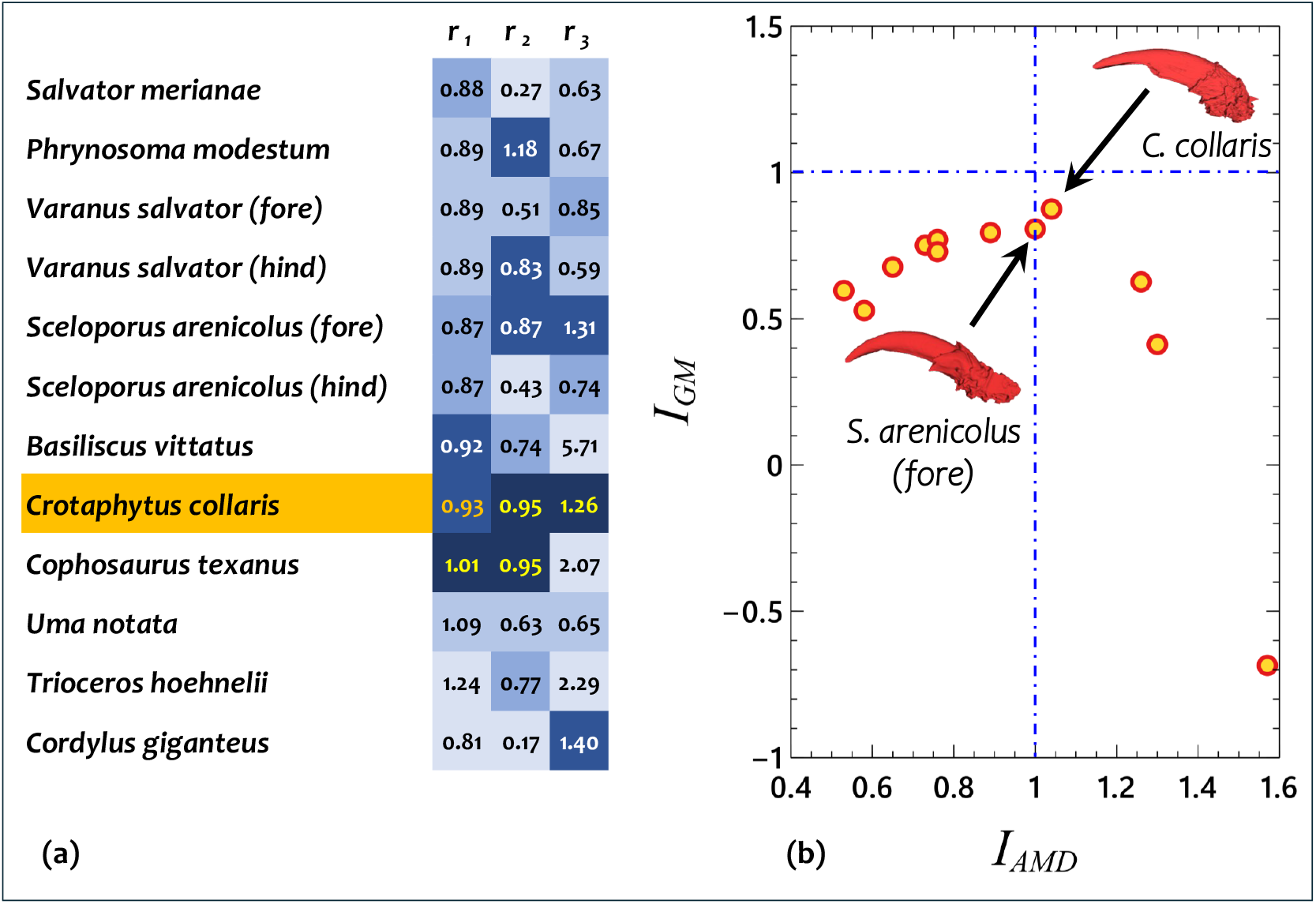
(a) choropleth map comparing *r*_1_, *r*_2_ and *r*_3_ for each of the species. The closest numbers to unity are highlighted in yellow and the next closest in orange. Boxes closest to unity are darkest while those farthest from unity are lightest. *C. collaris* is highlighted as it has the highest number of instances where it is close to 1. (b) Index *I*_*GM*_ plotted against index *I*_*AMD*_ for each of the species. The species closest to unity (identified in each axis by broken blue lines) are noted on the map.

The values closest to 1 are the superior claw geometries for use as our end effector. We note that while *C. texanus* scores exceptionally well in relation to the metrics *r*_1_ and *r*_2_, *C. collaris* scores well in all three of the metrics. The superiority of the *C. collaris* geometry is further reinforced by Figure 5(b), where the indices *I*_*GM*_ and *I*_*AMD*_ are plotted against one another. In this graph, the claws occupying positions closest to unity (shown as dotted blue lines) for both indices fulfill our design requirements for a claw geometry. As can be observed, there are two claws that are close to unity in both *I*_*GM*_ and *I*_*AMD*_, *S. arenicolus* and *C. collaris*. Since *C. collaris* shows itself as having optimal geometric qualities when comparing metrics *r*_1_-*r*_3_, as well as with respect to both *I*_*GM*_ and *I*_*AMD*_, we deduce that *C. collaris* is the obvious choice of claw for the robotic manipulator.

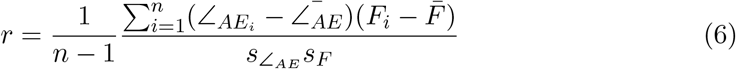

## 3. Discussion

The work described herein aimed to select an optimal claw type from amongst several claw types, one which is best suited for application to a simple bioinspired robotic manipulator. The gripping efficiency of lizard claws as previously researched [4], indicates that morphological parameters tend to function collectively as opposed to individually in the optimisation of claw grip efficiency. Here, we identify specific geometrical ratios that we deem are important collective parameters that contribute to the grip efficiency of a claw to a ping pong ball. Included is a ratio, *r*_1_ (cf. Equation 1), which measures the angle of the robot arm relative to the tip of the claw (end effector), against an idealised angle for a ball ∠_*AE*_, which we note is negatively correlated to force (cf. Figure 4). Also included is a ratio, *r*_2_ (cf. Equation 2), which measures the closest circular inner radius of the claw under scrutiny, against the radius of the ping pong ball. The importance of radius in a robotic manipulator comprising claws is well known, [31,32]. Rather than solely focus on claw radius as a lone concept, we normalise it against that of the object being gripped as it better contextualises gripper efficiency as discussed by Zhao et al. [33]. This is important in hard grippers like claws, as in soft grippers, the radius can be reached through material compliance [34] rather than through compatibility in meshing. Finally, the third ratio, *r*_3_ (cf. Equation 3), measures the maximum vertical distance travelled by the claw during testing, against a travel distance to the point where slip should commence, which would be half the ball radius, 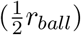. Slip occurs when the applied tangential force exceeds the maximum static friction force. The tangential forces increase with distance from the grip point from the south pole (the contact interface with the claw) and these forces decrease at approximately 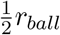 due to the shape of the claw and its relative position to the ball (it reaches a maximum ‘uphill’ climb). Slip therefore occurs on a sphere much earlier and more abruptly than in other primitives such as cuboids or cylinders [35], and is additionally a material dependent property [36,37]. Based on the choropleth map shown in Figure 5, we deduce that the claw from *C*.*collaris* most closely aligns with our optimal design requirements, as it has either the closest or second closest values to unity in each of the three metrics.

While the three metrics, *r*_1_-*r*_3_, individually provide a good comparative basis from which we can select a claw type, the indices, *I*_*GM*_ and *I*_*AMD*_, are beneficial as they form conjugates of all three. On comparison, we note that both *S. arenicolus (fore)* and *C. collaris* are very good candidates as they both have indices close to unity. Since of the two, *C. collaris* also scored well in the individual metrics (*r*_1_-*r*_3_), we select the claw geometry of *C. collaris* as the end effector for our robotic manipulator. Our selection methods focussed on gripping a ping pong ball as it is a challenging object to grip with a small size, light weight, and smooth surface, making precise gripping and manipulation difficult [30,38]. Claw inspired end effectors have received increasing attention in recent times as they have versatility in the types of structures they can grip [39–41]. We demonstrate a robotic grasp mechanism on a ping pong ball in Figure 6. Here, the ball is gripped by the tips of the claws, as the position and lightweightness of the ball prevents the tool from successfully performing a scooping motion. The ball is nevertheless held due to the tangential forces exerted by the mechanism causing sufficient friction at the claw tips. Since the ball is a 3-dimensional object, the contact force vector, *f*_*i*_, at the tip of each claw can be defined by Equation 7 [42], where the tangential force components are represented by *c*_1_ and *c*_2_, while the *c*_3_ is the normal force component. The component *c*_4_ represents the normal torque component, which will naturally be present in a grasped 3-dimensional object, the friction cone, *FC*_*i*_, for which is represented by Equation 8. In this equation, *ζ >* 0 represents the rotational static friction coefficient, *µ* is the translational static friction coefficient, *f*_*i*_ represent the contact forces vector at a point, *f*_3_ is the third individual force vector, and ℝ^3^ defines the set of all 3-dimensional real vectors.

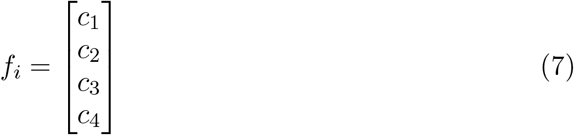

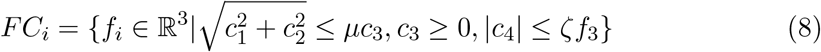

**Figure 6.**
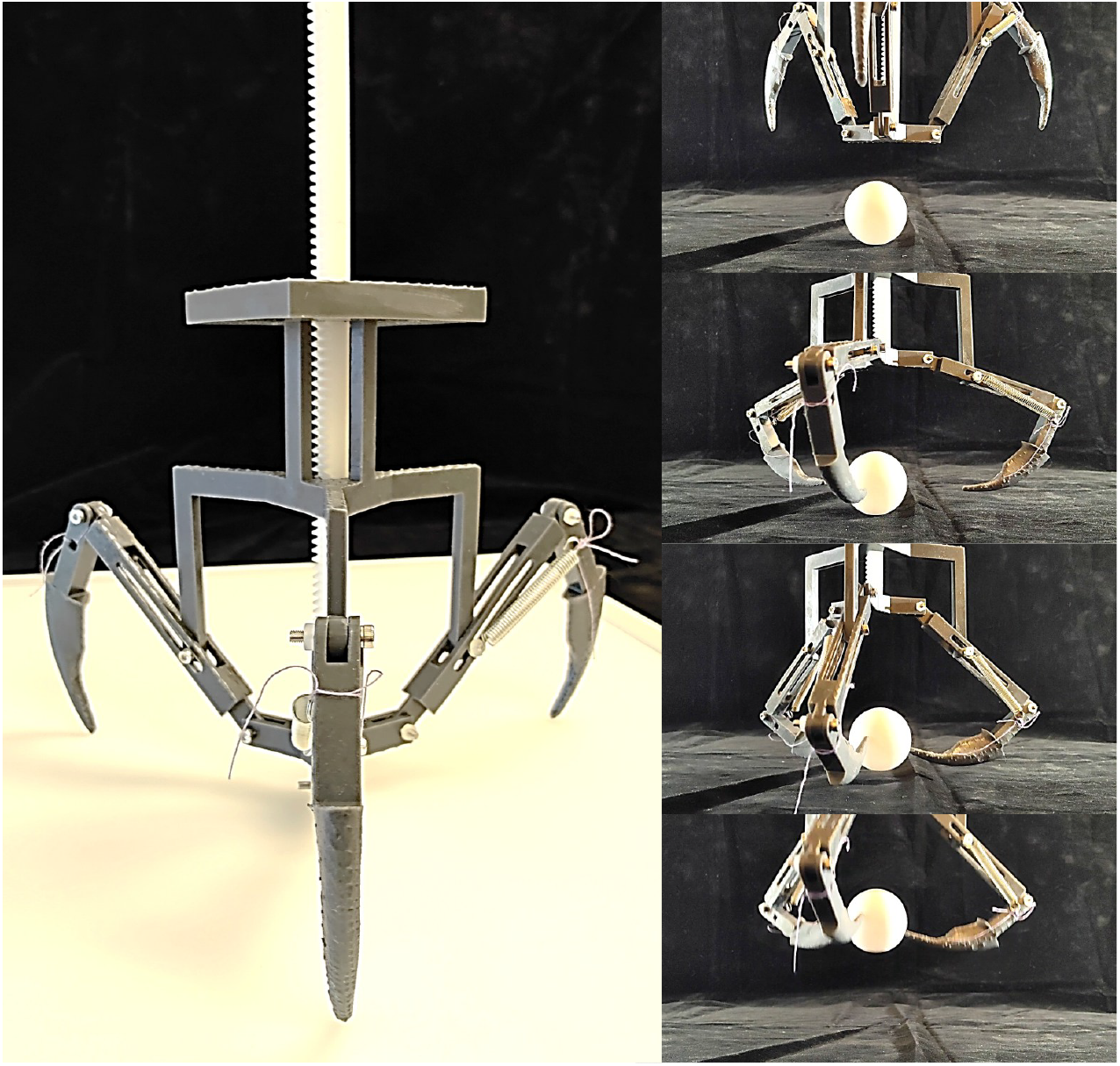
Finalised prototype (left) and a sequence of the gripper grasping and lifting a ball (right).

The demonstration of grasp using a scooping motion is easier in larger items. These in turn make use of a larger geometrical section of each claw than the ping pong ball example provided earlier. We evidence grasping after scooping in the sequence shots of Figure 7, where we grasp (a) an abalone shell (b) a boxing mitt and (c) an empty spool. The scooping motion is useful in a robotic manipulator as it enables a robot to effectively acquire and secure objects that are difficult to grasp with simple pinch or suction methods [23,28,43]. Scooping has the advantage of allowing the robot to push or slide under objects from a surface and stabilise and contain objects within the end effector (claw). Each of these enables a robot manipulator to work more effectively within certain environments, e.g. cluttered environments, where the ability to tilt into a grasp would be beneficial. This motion is especially beneficial in applications where the robot scoops fragile objects as grip can be adapted in real time based on the angle and style of grasp, effectively avoiding breakage whilst maximising acquisition effectiveness [44–46]. When our robotic manipulator scoops to grasp, it utilises the full benefit of the claw end effectors, which cradle the object (cf. Figure 7). While this is perhaps the most stable grasp, our manipulator is also able to grasp by pinching in order to grip smaller, lighter objects such as ping pong balls (cf. Figure 6). The dual ability to grasp through scooping, and to grasp through pinching is an important characteristic of our manipulator, as pinching allows for the careful and precise handling of objects of various shapes and fragility levels [46–48]. The main differentiation between the two rests on the careful control of the kinetics of digit motion.

**Figure 7.**
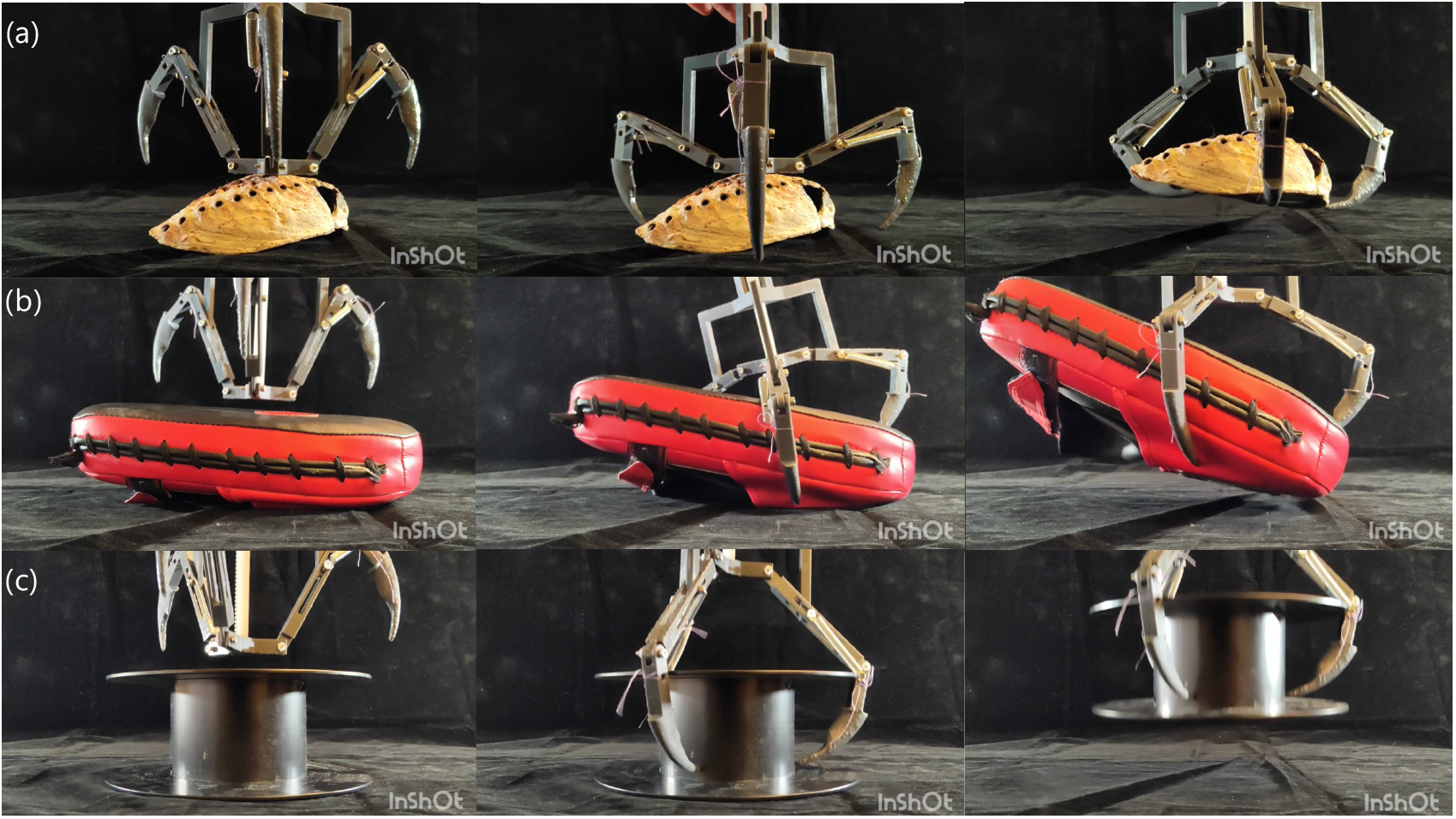
Using the prototype grasping tool to pick up (a) an abalone shell (b) a boxing mitt and (c) an empty spool (sequence shots are shown).

## 4. Conclusions

The work presented here identifies and selects an optimal gripping claw type for application to a bioinspired robotic manipulator, drawing on insights from lizard claw morphology coupled to grip efficiency. By focussing on three key geometrical ratios (i) arm to claw tip angle relative to an idealised angle, (ii) claw inner radius normalised against the object radius, (iii) and claw vertical travel distance relative to a slip distance threshold, our work provides a comprehensive framework for the evaluation of claw grip performance using a challenging object (ping pong ball) as a test archetype. Our final selection of the claw geometry from *C. collaris* is supported by both the individual metrics described above, as well as composite indices that conjoin them.

The full suite of metrics highlight the importance of using collective morphological parameters to design for stable and effective grip in robotics. Our robotic manipulator showcases versatility through its dual ability to perform pinching grasps on lightweight small objects, as well as scooping grasps on larger, irregular shaped objects. Our research advances the design principles needed for robust, bioinspired robotic claws as end effectors that integrate biomechanical insights into engineered gripping.

## 5. Methods

### 5.1. Claw procurement, selection, pretreatment and scanning

Claws were collected from adult specimens in the teaching collection of the Biodiversity Research and Teaching Collection at Texas A&M University. Species were chosen to include a diversity of ecologies and locomotor modes, and based on availability for destructive sampling. For each specimen, the longest digit of the forelimb and hindlimb was removed at or just proximal to the distal interphalangeal joint and placed in 70% ethanol for storage. Claw shape was quantified using micro-computed tomography (CT). Claws were scanned using a Bruker Skyscan 1273 in high resolution, from 3.8 to 10 m, 40 kV, and 85 to 100 A. Reconstructions were segmented in 3D Slicer [49] to isolate the claw. Volumetric data were converted into surface meshes and exported as.obj files for further analysis.

### 5.2. Computer aided design (CAD)

The generation of a CAD model of the claw from the CT scans required an initial qualitative assessment of the shape of the claw, to determine the parts of the claw that were exposed, relative to those that were underlying skin tissue. Key features of the claw were differentiated from scanning artefacts by visual analysis of the CT scan outputs. Often, the boundary between exposed claw and bone was clear, with the exposed claw typically showing a smooth curvature, while the skin or bone material typically exhibited no curvature, or showed topographical irregularities. This interface between the two materials was taken as the cut-off point for claw isolation, as applied to the grip test specimens. To simplify claw structures, we removed loose vertices and remeshed the surface of the claw using larger triangulation in Blender (Blender Foundation, Amsterdam, Netherlands), an open-source modelling and rendering software. The.stl file output from Blender (Blender Foundation, Amsterdam, Netherlands) was then imported into SolidWorks (Dassault Systmes SolidWorks Corporation, Waltham, Massachusetts, USA) and converted into a solid body to design-in features for grip testing. In these grip test specimens, the skin and bone sections of the claw were removed and scaled, such that the width of the claw at its widest point was 10 mm. A shaft was extruded from the cut segment of the bone. The extruded portion comprised a 7 *×* 7 *×* 100 mm shaft, which was extruded in line with the original direction of the bone. This was to ensure that loading aligned with the natural position of the claw. The claw and shaft model was then exported as an.stl for Fused Deposition Modelling (FDM) printing. Figure 8 illustrates the difference between an original scan, the smoothed mesh, and the final cleaned claw as attached to the attachment.

**Figure 8.**
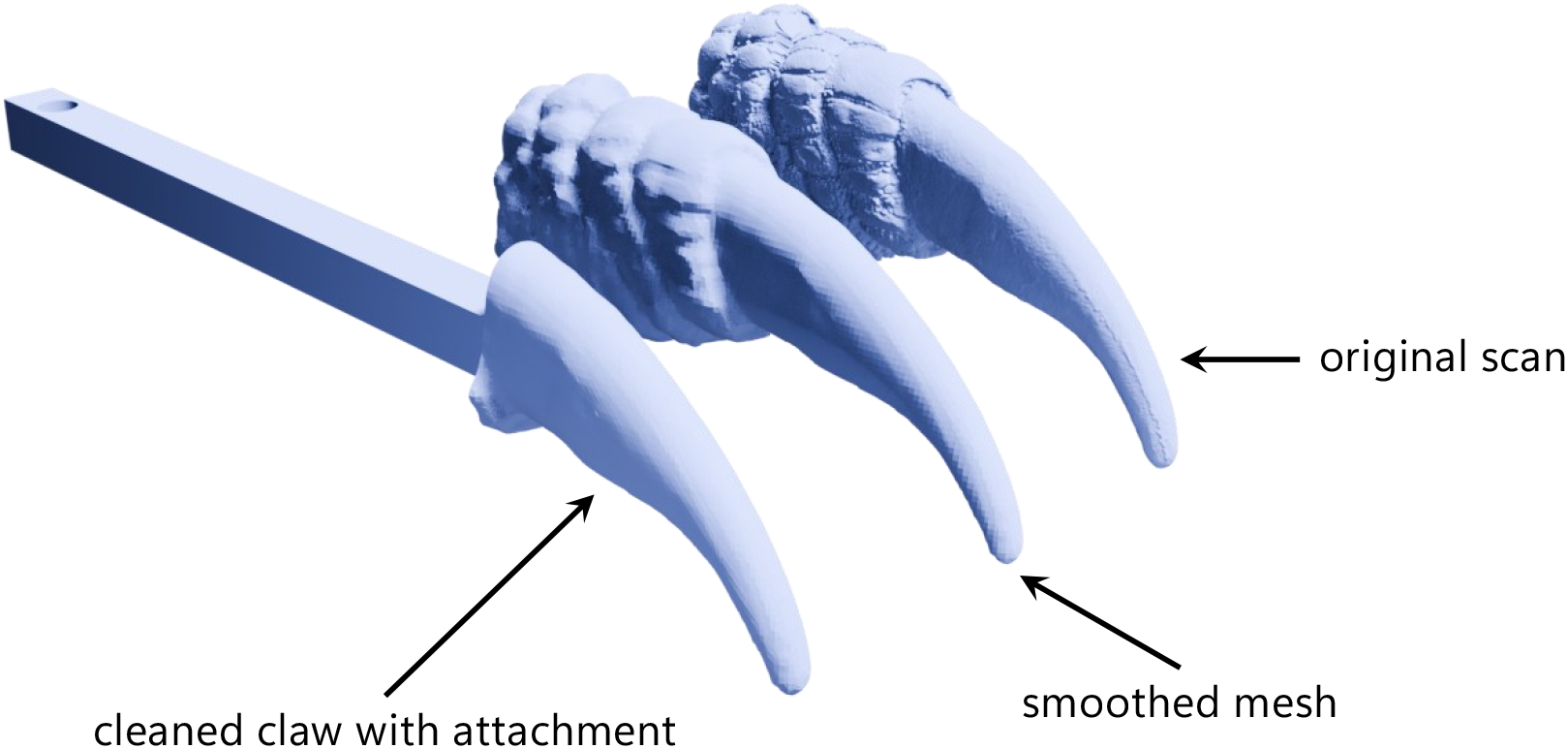
Comparison between original scan, smoothed mesh, and cleaned claw with attachment.

### 5.3. Manufacture

FDM slicing software Bambu Studio (Bambu Lab, Shenzhen, China) was used to generate G-code for a Bambu A1 Mini (Bambu Lab, Shenzhen, China) FDM printer. We printed the parts using a 15% infill, a grid infill pattern, and tree supports for prints oriented with the print bed parallel to the parasagittal plane of each claw with attachment. Two holes were drilled through a ping pong ball to stabilise it for testing. One hole was drilled at the ball’s north pole to fit both the bolt shaft and head, and one at the south pole, which only allowed a shaft to pass through. This maximises the area of the ping pong ball that can be gripped onto during the grip test, whilst concurrently securing the ping pong ball throughout the duration of the test.

The robotic manipulator was manufactured with using a Form 4 (Formlabs, Inc., Somerville, Massachusetts, USA) SLA printer as this provided excellent dimensional accuracy. The dedicated slicing software for Form 4 (Formlabs, Inc., Somerville, Massachusetts, USA) automatically generates the support and slices the.stl files. M3 bolts and nuts were used to connect joints during assembly. Springs were used between the second and the third digit of the manipulator to provide natural tension during grasping actions, and these were also secured using M3 bolts. A toothed rack was used to guide the first digit in the manipulator.

### 5.4. Grip testing

A low crosshead displacement rate of 5 mm/min was used during grip testing to ensure quasi-static behaviour of assembly. The arm base (also 3D-printed) held three claw attachements at inter-attachment angles of 120^°^ from centroid to centroid. These were attached using a brass insert and M5 bolts. The ping pong ball was attached to the fixture with an M12 bolt.

### 5.5. Design of robotic manipulator

Mimicry of grasping motion from digitisation, while ensuring that the claws could move simultaneously from single source was a considerable design challenge. The central mount interfaces with a rack and acts as a lever for the first digit. In the first digit, the routed slot was designed to allow a trajectory that draws the three first digits to the centre, while maintaining a simple geometry. The second digit has a spring connected to the first digit, and this is used to restore its position to a closed position in relation to the first digit. The claw from *C. collaris* was chosen as the end effector to apply to the robotic grasping tool. Figure 9 shows CAD renders of the final grasping tool including separate renders of the central rack mechanism, and the interdigit routed slot mechanism.

**Figure 9.**
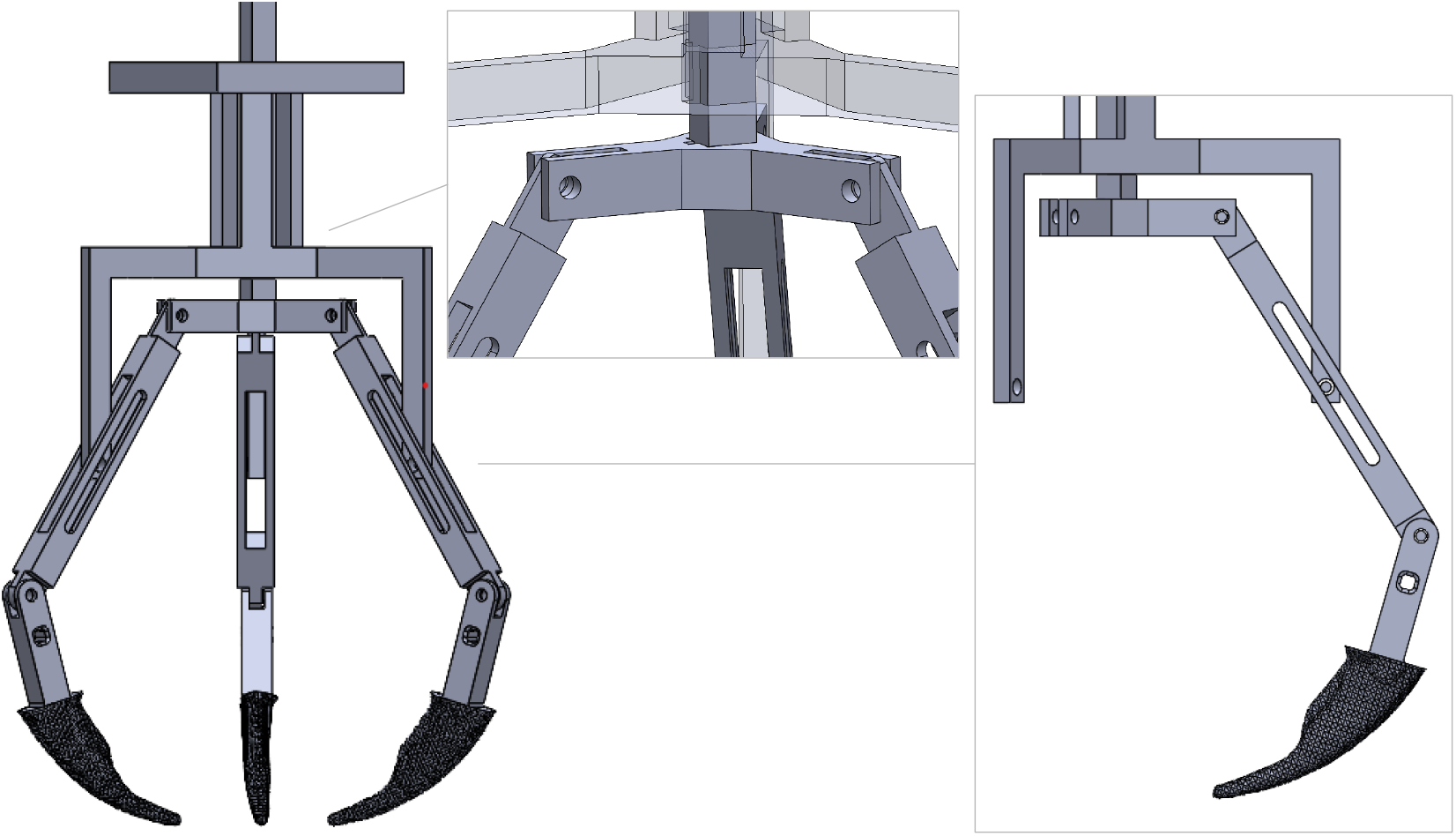
CAD renders of the final grasping tool including separate renders of the central rack mechanism, and the interdigit routed slot mechanism.

## Supporting information

Electronic Supplementary Material

## 6. Electronic Supplementary Materials

Maybe the tables…

## 7. Data Availability

1. Electronic Supplementary Tables (forces and displacements for all claw types)
2. Supplementary Video 1 - Ping Pong Ball
3. Supplementary Video 2 - Abalone Shell
4. Supplementary Video 3 - Boxing Mitt
5. Supplementary Video 4 - Spool

## 8. Acknowledgments

Edinburgh-Rice Strategic Collaboration Awards.

## 9. Author Contributions

Conceptualization (PA); Data curation (HL, KMD, AB); Formal analysis (PA); Funding acquisition (PA, AR); Investigation (HL); Methodology (HL, PA, KMD, AB, AR); Project administration (AR, PA); Resources (PA, AR); Software (-); Supervision (PA, AR); Validation (HL, PA, AR); Visualisation (PA, HL); Roles/Writing - original draft (PA, HL, AR); Writing - review and editing (AR, PA).

