## Supplementary material for "Grip and grasp: lizard claw inspired robotic manipulators": Electronic Supplementary Material: Electronic Supplementary Tables.pdf

ELECTRONIC SUPPLEMTARY MATERIAL

### Grip-grasp efficiency of bioinspired robotic lizard claws

**ARTICLE HISTORY**
Compiled October 22, 2025

#### 1. Property Tables

**Table 1.** Grip force of various claw types. STD - standard deviation, SE - standard error, CoV - coefficient of variation.

| Species | Value [N] | STD [N] | SE [N] | CoV [%] |
|---|---|---|---|---|
| *Salvator merianae*(hind) | 2.6 | 0.8 | 0.9 | 29.8 |
| *Phrynosoma modestum* (fore) | 6.7 | 2.2 | 2.4 | 32.7 |
| *Varanus salvator* (fore) | 3.8 | 1.2 | 1.3 | 30.6 |
| *Varanus salvator* (hind) | 3 | 0.4 | 1.1 | 14.2 |
| *Sceloporus arenicolus* (fore) | 12.5 | 3.2 | 4.4 | 25.3 |
| *Sceloporus arenicolus* (hind) | 7.4 | 3.1 | 2.6 | 41.9 |
| *Basiliscus vittatus* (hind) | 8.7 | 2.1 | 3.1 | 24 |
| *Crotaphytus collaris*(fore) | 12.9 | 4.6 | 4.6 | 35.9 |
| *Cophosaurus texanus* (fore) | 13 | 7.5 | 4.6 | 57.9 |
| *Uma notata* (hind) | 19.8 | 17.5 | 7 | 88.5 |
| *Trioceros hoehnelii* (hind) | 16.6 | 10.6 | 5.9 | 63.5 |
| *Cordylus giganteus* (hind) | 0.1 | 0.009 | 0.04 | 8.7 |

**Table 2.** Grip displacement of various claw types. STD - standard deviation, SE - standard error, CoV - coefficient of variation.

| **Species** | **Value [%]** | **STD [%]** | **SE [N]** | **CoV [%]** |
|---|---|---|---|---|
| *Salvator merianae* (hind) | 79 | 27.5 | 27.9 | 34.8 |
| *Phrynosoma modestum* (fore) | 75.4 | 33 | 26.7 | 43.7 |
| *Varanus salvator* (fore) | 58.6 | 28 | 20.7 | 47.9 |
| *Varanus salvator* (hind) | 84.7 | 20.7 | 30 | 24.4 |
| *Sceloporus arenicolus* (fore) | 38.1 | 10.5 | 13.5 | 27.5 |
| *Sceloporus arenicolus* (hind) | 67.5 | 19.6 | 23.9 | 29.1 |
| *Basiliscus vittatus* (hind) | 8.8 | 5.1 | 3.1 | 58.6 |
| *Crotaphytus collaris* (fore) | 39.8 | 18.3 | 14.1 | 46 |
| *Cophosaurus texanus* (fore) | 24.2 | 8.6 | 8.6 | 35.4 |
| *Uma notata* (hind) | 76.9 | 40.2 | 27.2 | 52.3 |
| *Trioceros hoehnelii* (hind) | 21.8 | 6.1 | 7.7 | 28.2 |
| *Cordylus giganteus* (hind) | 35.8 | 9.4 | 12.6 | 26.2 |
